# Posttranslational modifications of the DUX4 protein impact toxic function

**DOI:** 10.1101/2022.07.22.501148

**Authors:** Renatta N. Knox, Jocelyn O. Eidahl, Lindsay Wallace, Sarah Choudury, Afrooz Rashnonejad, Nizar Y. Saad, Michael E. Hoover, Liwen Zhang, Owen E. Branson, Michael A. Freitas, Scott Q. Harper

**Affiliations:** Center for Gene Therapy, The Abigail Wexner Research Institute at Nationwide Children’s Hospital, Columbus, OH 43205, USA; Department of Pediatrics, The Ohio State University College of Medicine, Columbus, OH 43210, USA; Department of Cancer Biology and Genetics, The Ohio State University College of Medicine, Columbus, OH 43210, USA

**Author notes:** These authors contributed equally to this work (Renatta N. Knox, Jocelyn O. Eidahl). To whom correspondence should be addressed: Scott Q. Harper; Center for Gene Therapy, The Research Institute at Nationwide Children’s Hospital, Columbus, OH 43205.

## Abstract

**Objective:** Facioscapulohumeral muscular dystrophy (FSHD) is caused by abnormal de-repression of the transcription factor DUX4, which is toxic to muscle *in vitro* and *in vivo*. While the transcriptional targets of DUX4 are known, the regulation of DUX4 protein and the molecular consequences of this regulation are unclear. Here, we used *in vitro* models of FSHD to identify and characterize DUX4 posttranslational modifications (PTMs) and their impact on the toxic function of DUX4.

**Methods:** DUX4 protein was immunoprecipitated and mass spectrometry performed to identify PTMs. We then extensively characterized DUX4 PTMs and potential enzyme modifiers using mutagenesis, proteomics and biochemical assays in human cell lines and human myoblast cell lines.

**Results:** Our *in vitro* screen of DUX4 PTM mutants identified arginine methyl-null and serine/threonine phosphomimetic mutants that protected cells against DUX4-mediated toxicity and reduced the ability of DUX4 to transactivate downstream gene targets, including FSHD biomarkers. Using additional proteomics and biochemical approaches, we identified protein kinase A (PKA) and a protein arginine methyltransferase (PRMT1) as components of the DUX4 complex. Importantly, over-expression of PRKACA, a catalytic subunit of the PKA holoenzyme, mitigated DUX4 toxicity, while pharmacologic inhibition of PRMT1 protected human myoblasts from DUX4-mediated apoptosis.

**Interpretation:** These results demonstrate that DUX4 is regulated by PTMs and that DUX4 PTMs, or associated modifying enzymes, may be druggable targets for FSHD therapy.

## INTRODUCTION

Facioscapulohumeral muscular dystrophy (FSHD) is among the most common forms of muscular dystrophy, affecting an estimated 870,000 people worldwide. FSHD is typically characterized by slowly progressive, asymmetric weakness affecting muscles of the face, shoulders, trunk and extremities. FSHD is associated with significant disability that may lead to wheelchair dependence ^1^. Most patients present in late adolescence and early adulthood, however rare early onset forms show greater disease severity and systemic manifestations ^2,3^. Unfortunately, there are no approved disease modifying treatments and care remains supportive.

In the simplest terms, FSHD is caused by abnormal de-repression of the *DUX4* gene in skeletal muscle ^4^. *DUX4* produces a double homeodomain transcription factor (DUX4) that normally operates in early embryogenesis and is repressed in adult tissues except the thymus, testes and possibly skin ^5^. When expressed in skeletal muscle, DUX4 activates numerous pathways that are incompatible with normal muscle function, including oxidative stress and apoptosis, among others ^6–11^. The *DUX4* open reading frame (ORF) is embedded within D4Z4 macrosatellite repeats located on the human chromosome 4q35. In non-FSHD muscle, this region is typically hypermethylated, embedded in heterochromatin and *DUX4* is not expressed. In FSHD, shorter D4Z4 arrays (FSHD1) or mutations in chromatin modifier genes (FSHD2) lead to 4q DNA hypomethylation, enabling transcription of *DUX4* ^4,12–15^. If this occurs on a permissive allele (4qA) containing a polyA signal for *DUX4*, the *DUX4* mRNA arising from the most distal telomeric D4Z4 repeat is polyadenylated, stabilized, and translated into toxic DUX4 protein ^4,6,7^

Since the discovery that *DUX4* is the causative gene in FSHD, there has been great interest in developing *DUX4* inhibition therapies. Since DUX4 protein activates toxic pathways, arguably the most effective *DUX4* inhibition strategies will silence *DUX4* at the DNA or RNA levels, thereby avoiding protein production. Indeed, several approaches to accomplish *DUX4* DNA or mRNA silencing have been previously described, including by our laboratory ^16,17^. These strategies, although promising, offer no remedy for inhibiting DUX4 after protein has already been produced. To date no one has yet described an approach to directly interfere with the ability of DUX4 protein to transactivate downstream genes. This was the overall goal of the current study.

To develop DUX4 protein inhibitors, it is critical to understand how DUX4 protein functions and is regulated. Previous studies have defined domains within the DUX4 protein that are required for DUX4 transactivation activity and toxicity, including two N-terminal DNA binding homeodomains, nuclear localization signals, and a C-terminal transactivation domain that recruits histone acetyltransferases (p300/CBP) to target genes ^7,18–21^. Once incorporated into a protein, primary amino acids can be modified at the post-translational level to regulate and alter protein function. These posttranslational modifications (PTMs), link gene regulation machinery with signal transduction pathways and are known to impact the function, stability, and subcellular localization of transcription factors ^22,23^. We hypothesized that the DUX4 transcription factor was regulated by PTMs and that prospective modifications and the enzymes that add or remove them could be targets for DUX4 inhibition strategies.

## METHODS

### Mutagenesis

*DUX4* modification mutant plasmids were constructed using a recombinant PCR method. Mutants were constructed using PCR to amplify the *DUX4* ORF, with primers containing the mutated site, using CMV driven wildtype *DUX4* plasmid (AAV.DUX4.V5) as PCR template. The entire mutant *DUX4* ORF was amplified, gel purified and cloned into PCR-blunt II-Topo, prior to sequence verification. *DUX4* mutant ORFs were then cloned into AAV.CMV.DUX4 or AAV.CMV.eGFP to replace either wild-type DUX4 or eGFP, respectively. Phospho-null, Phosphomimic, Methyl-null, basic and Methyl-mimic mutants were synthesized by Genscript with N-terminal NheI and C-terminal Acc65I restriction enzyme site s flanking the *DUX4* ORF, and then subcloned into AAV.CMV.eGFP.

### Cell Culture

Human embryonic kidney cells (HEK293) and human immortalized myoblasts (WS236, 15V, biceps, unaffected control cells, iDUX4 cells) were maintained as previously described ^24,25^. Briefly, HEK293 cells were cultured in DMEM supplemented with 10% fetal bovine serum, L-glutamine and penicillin/streptomycin at 37°C in 5% CO_2_. Human immortalized myoblasts were cultured in LHCN media containing DMEM supplemented with 16% Medium 199, 15% fetal bovine serum,30⍰ng/ml zinc sulfate,1.4 µg/ml vitamin B12,55⍰ng/ml dexamethasone,2.5⍰ng/ml human growth factor,10 ng/ml fibroblast growth factor, 20⍰mM HEPES and penicillin/streptomycin. iDUX4 human immortalized myoblasts were cultured in F10 medium supplemented with 20% fetal bovine serum, 10ng bFGF,1µM dexamethasone and penicillin/streptomycin.

### Protein Immunoprecipitation

HEK293 cells were transfected with a total of 4 µg AAV.CMV.DUX4.V5, PRMT1-GFP, or PRKACA-Flag (1×10^6^ cells/well) using Lipofectamine 2000 and harvested 16 hours later using cold 1 x PBS. Cells were pelleted and lysed in Buffer A containing 137 mM NaCl, 50 mM Tris, pH 7.5, 1 % NP-40, protease inhibitor cocktail (Sigma), and phosphatase inhibitor (Roche). All steps were performed at 4°C or on ice. Lysate was incubated with protein agarose G for 1 hour, while rotating. The supernatant was subsequently incubated with anti-V5 antibody conjugated to agarose resin overnight, while rotating. The resin was washed five times with Buffer A, then resuspended in Buffer A supplemented with 1 mM DTT and 1 x LDS-PAGE sample buffer (Invitrogen). DUX4 complexes were eluted by boiling at 95°C for 10 min.

### Liquid Chromatography and Mass Spectrometry (LC-MS/MS)

Immunoprecipitated samples were loaded into a TGX 4-15 % precast gel (Bio-Rad), resolved and stained with Bio-Safe Coomassie (Bio-Rad). The DUX4 band was excised and disulfide bonds reduced and alkylated. Bands digested overnigt at 37 °C with 800 ng of trypsin (Promega) and/or chymotrypsin (Promega) in 100 mM ammonium bicarbonate (Sigma). Peptides were extracted from the gel, dried and resuspended in loading buffer (2% acetonitrile, 0.1 % formic acid). Peptides were separated on a Thermo Dionex UltiMate 3000 RSLC HPLC coupled to a Thermo Orbitrap Fusion Tribrid mass spectrometer. Peptides were separated on a PepMap100 C18 microcolumn (5 µm,100Å,0.3×50mm) using a linear gradient 5-30% of acetonitrile in water with 0.1% formic acid. MS/MS data were collected using data dependent acquisition mode.

### Mass Spectrometry Data Analysis

RAW data were converted to the mzML format using the MSConvert tool in ProteoWizard (v3.0.4624) and searched against a database containing the DUX4 sequence (UniProt accession Q9UBX2) with a C-terminal V5 epitope tag (GKPIPNPLLGLDST) and common contaminant proteins (downloaded 22 June 2015; 234 total entries) using the MassMatrix search engine v2.4.2 and MASCOT (version 2.6.0, Matrix Science). Peptide mass tolerance was set at 20 ppm with a fragment mass tolerance of 0.02 Da. against UniProt human database containing the DUX4 sequence (UniProt accession Q9UBX2) with a C-terminal V5 epitope tag (GKPIPNPLLGLDST). Peptide mass tolerance was set at 10 ppm with a fragment mass tolerance of 0.05 Da. Variable modifications included acetylation of K; mono-, di- or tri-methylation of K; mono- or di-methylation of R; oxidation of M; and phosphorylation of S,T,Y. Carbamidomethylation of C was included as a fixed modification. Enzyme specificity was set for trypsin and chymotrypsin with up to four missed cleavages.

### Quantitative PCR

HEK293 cells and human myoblasts (5×10^5^ cells/well) were transfected with 2ug of AAV.DUX4.V5 (wild type or mutant construct) using Lipofectamine 2000 (Thermo Scientific). Cells were harvested 24 hours post-transfection in TRIzol RNA isolation reagent (Life Technologies). RNA was isolated, DNase treated and reverse transcribed into cDNA using the High-Capacity cDNA Reverse Transcription Kit (Applied Biosystems). TaqMan gene expression assays (Applied Biosystems) were used to quantify human *RPL13A* (Hs01494366_g1), human *ZSCAN4* (Hs00537549_m1) and human *PRAMEF12* (Hs04193637_mH). Efficiencies were comparable among all probes. *RPL13A* was used as the reference gene for normalization. The normalized expression (ΔΔCq) was calculated relative to control pClNeo transfected cells.

### Immunofluorescence

HEK293 cells (187,500 cells/well) were transfected in suspension with 800 ng AAV.DUX4.V5, AAV.DUX4 mutant constructs, or pCINeo (transfection control plasmid) using Lipofectamine 2000 reagent (Thermo Scientific). Cells were fixed using 4% paraformaldehyde (PFA) 20 hours post-transfection, then permeabilized with 0.25 % Triton X-100 in 1x PBS. Cells were then blocked using 3% bovine serum albumin (BSA) for 1 hour and incubated with anti-V5 primary antibody (1:2000; Abcam) in 1x PBS and 10 % normal goat serum overnight at 4°C. Next, the cells were incubated with Alexa-594 coupled goat anti-rabbit immunoglobulin G (IgG) secondary antibodies (1:5000; Molecular Probes) in 1x PBS for 2 hours at room temperature. All slides were mounted in Vectashield (Vector Laboratories) with DAPI.

### Western blot protein visualization

HEK293 cells (65,000 cells/well) were transfected in suspension with 100 ng AAV.DUX4.V5, AAV.DUX4 mutant constructs.V5 or pCINeo (transfection control plasmid) using Lipofectamine 2000 reagent (Thermo Scientific). Cells were harvested 24 hours post-transfection in RIPA buffer containing 10 mM Tris-Cl (pH 8.0), 1% Triton X-100, 0.1% sodium deoxycholate, 1% SDS, 140 mM NaCl and 1 mM DTT. Total protein was quantified using the Lowry Protein Assay (Bio-Rad) and analyzed by 12% SDS-PAGE. Protein was visualized by western blot using an anti-V5-HRP antibody (Invitrogen).

### *In vitro* Kinase Screen

A radiometric protein kinase filter-binding assay was used for measuring the kinase activity of the 245 protein kinases with recombinant DUX4 protein (Reaction Biology). The reaction cocktails containing kinase solution and buffer/ATP/test sample mixture, were pipetted into 96 well, V-shaped polypropylene microtiter plates. The reaction cocktails contained 60 mM HEPES-NaOH, pH 7.5, 3 mM MnCl_2_, 3 μM Na-orthovanadate, 1.2 mM DTT, 50 µg/ml PEG_2000_, 1μM ATP/[γ-^33^P]-ATP, protein kinase (1-400 ng/50μl) and DUX4 protein (5 µg/50 µl) with some reactions supplemented with CaCl_2_, EDTA, phosphatidylserine, 1.2-Dioleyl-glycerol, cGMP or calmodulin as needed. The assay plates were incubated at 30°C for 60 minutes. The reaction cocktails were stopped with 10 % (v/v) H_3_PO_4,_, transferred into 96-well glass-filter plates (MSFC, Millipore), pre-wetted with 150 mM H_3_PO_4_, followed by 30 min incubation at room temperature. The filter plates were washed three times with 150 mM H_3_PO_4_ and once with 100% ethanol. After drying the plates for 30 min at 40 °C, 50 μl of scintillator (Rotiszint Eco plus, Roth) were added to each well and incorporation of ^33^Pi and quantification of incorporated cpm was *determined* with a microplate scintillation counter (Microbeta, Perkin Elmer).

### Caspase-3/7 Activation Assay

HEK293 cells were transfected with 100 ng plasmid DNA (65,000 cells/well) using Lipofectamine 2000 and assays were performed 48 hours later using the Apo-ONE Homogeneous Caspase-3/7 Assay (Promega). Briefly, 100 µl of the reagent was added to each well and the 96-well plate was gently rotated for 20 min. Relative fluorescence was monitored every hour for 4-6 hours. Inhibitor studies were performed in human myoblasts which were transfected with 100ng of DUX4 DNA (65,000 cells/well) with Lipofectamine 2000 or iDUX4 myoblasts (40,000 cells/well) were induced with 1ug/mL of doxycycline (Fischer). The inhibitors adenosine dialdehyde (AdOx; Sigma), arginine methyltransferase inhibitor, (AMI-1; EMD Millipore), or salvianolic acid A (SAA; Sigma and AK scientific) were added 1.5 hours later and the caspase assay was performed 48 hours later as above.

### Cell viability assay

Human myoblasts (WS236, 15V, biceps, unaffected control cells) were transfected with DUX4 modification mutants using the Human Dermal Fibroblast Nucleofector kit (Amaxa). Cells were washed once with 1xPBS prior to trypsinization. Trypan blue was added (1:1) to the cell suspension. Total and trypan blue stained cells were counted using a hemocytometer. Cell viability was calculated using the following equation: [(1-(trypan blue stained cells/total amount of cells)) x100].

### DUX4-activated reporter in cells

HEK293 cells (65,000 cells/well) were transfected with 100 ng plasmid DNA (AAV.CMV.DUX4.V5 or mutants) and 100 ng pLenti.DUX4-activated GFP, previously reported in ^6^, using Lipofectamine 2000 (Thermo Scientific) and plated simultaneously on 96-well plates. GFP expression was quantified 24- and 48-hours post-transfection using the SPECTRAmax M2 instrument (Molecular Devices). GFP expression was visually monitored with a fluorescent stereo microscope (Leica M165 FC microscope, Leica Microsystems).

### Rapid Immunoprecipitation Mass Spectrometry of Endogenous Proteins (RIME)

The RIME assay was performed as previously described ^26^. Briefly, cells were fixed with 1% formaldehyde for 8 min and quenched with 0.125 M glycine. Chromatin was isolated by the addition of lysis buffer, followed by disruption with a Dounce homogenizer. Lysates were sonicated and the DNA sheared to an average length of 300-500 base-pairs. Genomic DNA (Input) was prepared by treating aliquots of chromatin with RNase, proteinase K and heat for de-crosslinking, followed by ethanol precipitation. Pellets were resuspended and the resulting DNA was quantified on a NanoDrop spectrophotometer. An aliquot of chromatin (100 µg) was pre-cleared with protein G agarose beads (Invitrogen). Proteins of interest were immunoprecipitated using 10 µg of anti-V5 (Abcam) and protein G magnetic beads. Protein complexes were washed, then trypsin was used to remove the immunoprecipitate from beads and digested the protein sample. Protein digests were separated from the beads and purified using a C18 spin column (Harvard Apparatus). The peptides were vacuum dried using a speedvac. Digested peptides were analyzed by LC-MS/MS on a Thermo Scientific Q Exactive Orbitrap Mass spectrometer in conjunction with a Proxeon Easy-nLC II HPLC (Thermo Scientific) and Proxeon nanospray source. All MS/MS samples were analyzed using X! Tandem (The GPM, thegpm.org; version CYCLONE (2013.02.01.1)). Scaffold (version Scaffold_4.6.1, Proteome Software) was used to validate MS/MS based peptide and protein identifications.

### Statistical Analysis

All statistical analyses (Apo-ONE Homoegeneous Caspase-3/7 Assay, Q-PCR, DUX4-activated GFP Reporter Assay) were performed in GraphPad Prism 5 (GraphPad Software, La Jolla, CA) using the indicated tests.

## RESULTS

To detect DUX4 post-translational modifications (PTMs) we transfected a V5-epitope tagged DUX4 expression plasmid in HEK293 cells, immunoprecipitated DUX4 with a V5 antibody and performed mass spectrometry (MS). We detected the DUX4 protein migrating near the expected molecular weight of 52kDa by SDS-Page (Fig 1A). Following in-gel digestion, we used high resolution mass spectrometry to detect modified DUX4 peptides. We used a trypsin or trypsin/chymotrypsin digestion strategy in two biological replicates and obtained 85% and 43% sequence coverage, respectively. We identified 17 prospectively modified amino acids, including novel DUX4 serine phosphorylation sites, arginine monomethylation and dimethylation sites, and a lysine acetylation site (Fig 1B, C and Table 1). We were unable to localize a subset of phosphorylation sites given their close proximity, however the spectral information indicated that these peptides were phosphorylated (Table 2).

**FIGURE 1.**
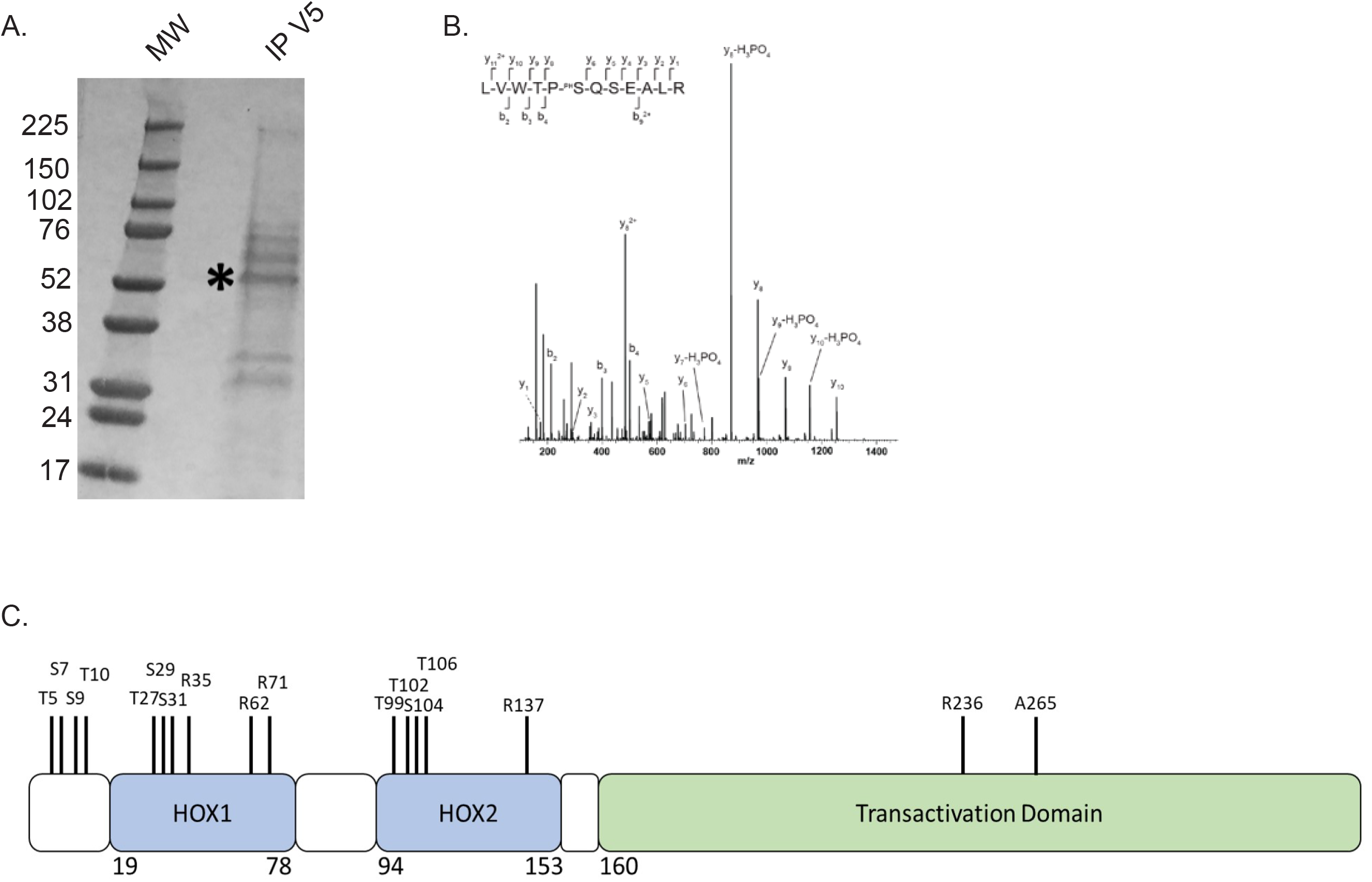
DUX4 undergoes post-translational modification in HEK293 cells. (A) Coomassie blue-stained SDS-polyacrylamide gel containing purified DUX4 protein. Asterisk (*) corresponds to 52 kDa, the molecular weight (MW) of the DUX4 protein. (B) Mass spectrum of the modified ‘LVWTPSQSEALR’ peptide. (C) Domain structure of DUX4 protein including homeodomain 1 (HOX1), homeodomain 2 (HOX2), and the C-terminal transactivation domain with the PTM sites identified by mass spectrometry.

**TABLE 1.**
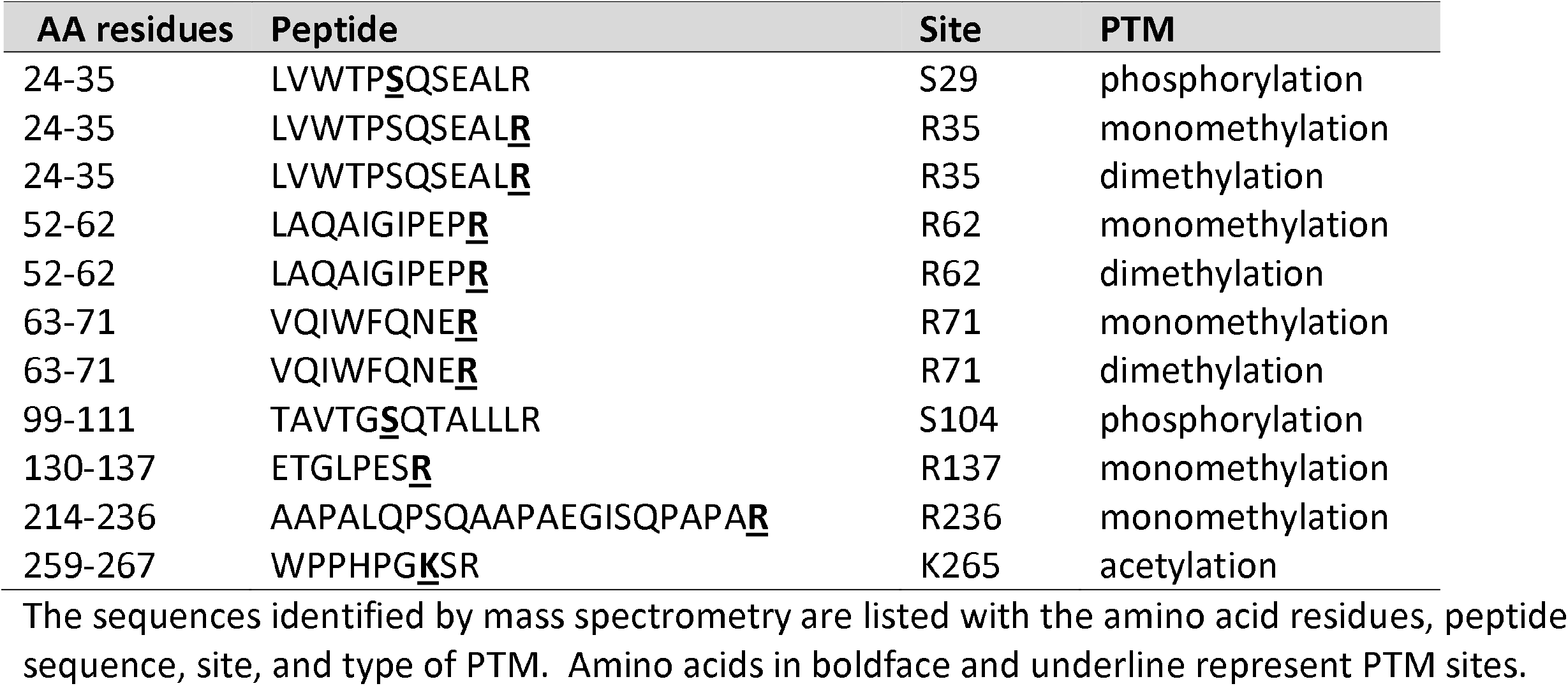
DUX4 peptides containing PTMs.

**TABLE 2.**
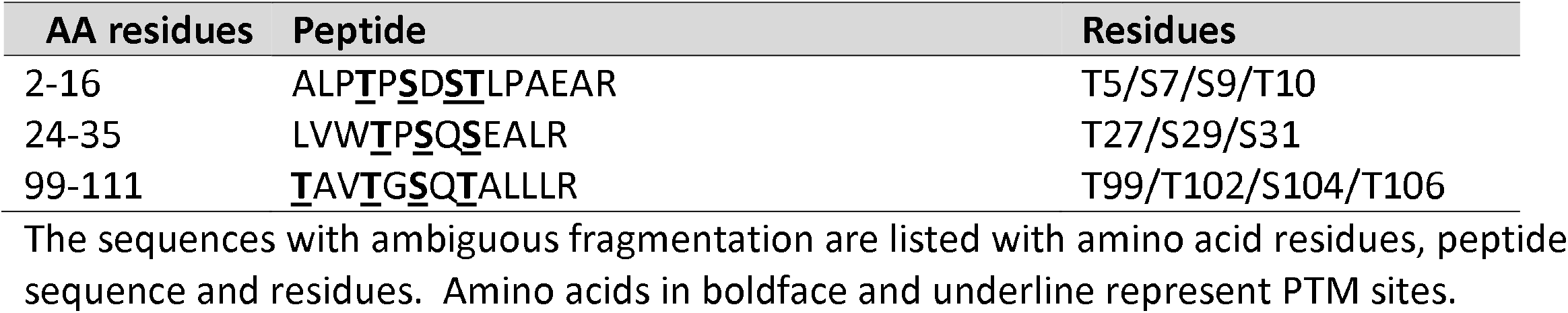
DUX4 peptides with ambiguous fragmentation.

To determine the functional consequence of prospective DUX4 PTMs, we generated 55 mutants designed to mimic or prevent the MS-identified phosphorylation, methylation, and acetylation sites (Supp Table 1). This total included DUX4 PTM mutants containing phosphorylation sites with ambiguous fragmentation. We generated 3 mutants for every methylated arginine residue and 2 for phosphorylated serine/threonine and acetylated lysine (Table 3). We also generated controls containing mutations in every putatively modified arginine or serine/threonine residue (DUX4-Methyl null and mimic; DUX4-Phospho null and mimic, respectively; Supp. Table 1).

**TABLE 3.** Mutagenesis strategy for DUX4 amino acids.

| Modified Amino Acid | Null Substitution | Mimic Substitution |
| --- | --- | --- |
| Methyl-Arg | 2 nulls per residue<br>Arg to Ala<br>Arg to Lys (maintains + charge) | 1 methyl mimic per residue<br>Arg to Leu |
| Phospho-Ser/Thr | 1 null per residue<br>Ser/Thr to Ala | 1 phospho mimic per residue<br>Ser/Thr to Glu or Asp |
| Acetyl-Lys | 1 null per residue<br>Lys to Ala | 1 acetyl mimic per residue<br>Lys to Gln |

Arg: arginine; Ser: serine; Thr: threonine; Lys: lysine; Ala: alanine; Leu: leucine; Gln: glutamine; Glu: glutamic acid; Asp: aspartic acid

Because DUX4 expression causes apoptosis in numerous cell types, we used a cell death assay as a primary screen to assess the functional impacts of DUX4 PTMs on DUX4-induced cell death ^7,9^. To do this, we transfected plasmids expressing wild-type DUX4 or DUX4 PTM mutants into HEK293 cells and performed caspase-3/7 apoptosis assays. We identified five mutants that significantly reduced DUX4-associated caspase-3/7 activation (Fig 2A-D). Among these 5 mutants, three contained different single mutations in the same methylated arginine residue (R71), such that methylation null and mimic mutations at R71 produced the same protective effect. The other 2 protective DUX4 mutants we identified in the caspase-3/7 screen contained phospho-mimic changes in 3 or 4 serine/threonine residues, respectively (DUX4-S29D/S31D/T106D; aka Mutant 2) and DUX4-T99E/T102E/S104E/T106E; aka Mutant 5) (Fig 2C). Interestingly, single mutations of prospective phosphorylated serine or threonine residues had no impact on DUX4 toxicity. Notably, the amino acids changed in the five protective DUX4 mutants were all located within the DUX4 homeobox DNA binding domains (Fig 1C).

**FIGURE 2.**
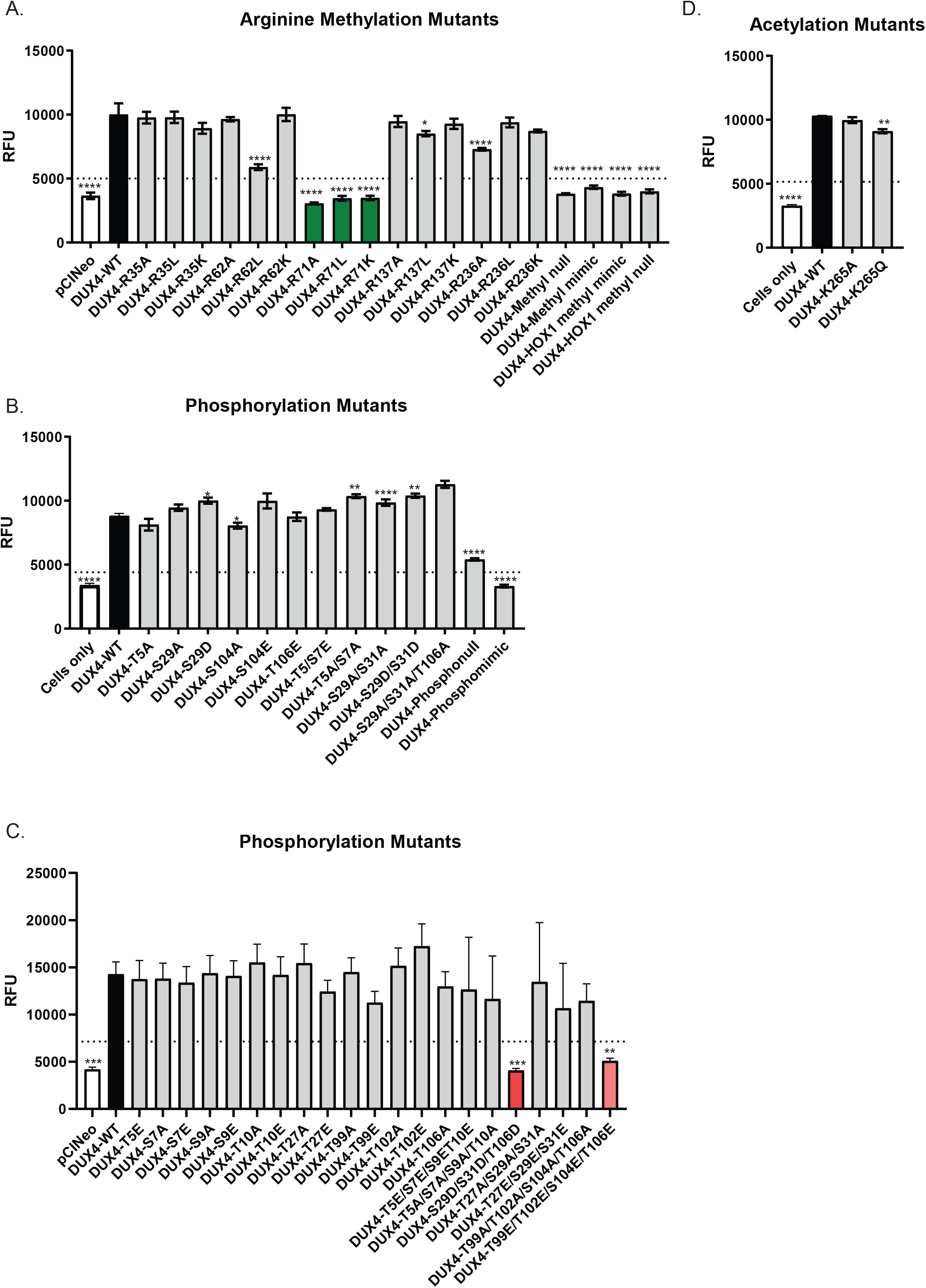
Identification of DUX4 PTM mutants that protect against apoptotic cell death. HEK293 cells were transfected with wild type DUX4 or the indicated DUX4 PTM mutant and the caspase 3/7 assay was performed 48 hours later to assess for cell death. Representative data for (A) arginine methylation mutants, (B, C) phosphorylation mutants and (D) acetylation mutants (D). Asterisk (*) indicates significant differences compared to DUX4-WT. * P<0.05, ** P<0.01, *** P<0.001, **** P<0.0001. One-way ANOVA test. Representative data for n=3-4 independent experiments performed in triplicate. Dashed line indicates the value which is 50% of DUX4-WT. RFU: relative fluorescence units.

DUX4 is a transcription factor that activates downstream target genes, and we therefore next sought to determine the impacts of DUX4 PTM mutations on its transactivation function. For this experiment we specifically focused on the 5 non-toxic mutants we identified in the caspase-3/7 assay. We first tested the ability of wild-type DUX4 or DUX4 PTM mutants to transactivate a previously described DUX4-responsive GFP reporter construct ^6^. While none of the 5 mutants were completely devoid of transactivation activity in this reporter assay, all failed to achieve GFP expression at the level produced by wild-type DUX4 (Fig 3 and 4). For the R71 mutants, we quantified a gradient of GFP expression (R71A<R71L<R71K; 26%, 32%, 59% of wild-type GFP levels) (Fig 4 A, B). Thus, maintaining a basic residue at this location (R71K) produced the most transactivation in this assay among the 3 R71 mutants. These data are consistent with our structural modeling suggesting the DUX4 R71 residue lies close to where DUX4 binds DNA and could be involved in making electrostatic interactions with DNA (Fig 4C).

**FIGURE 3.**
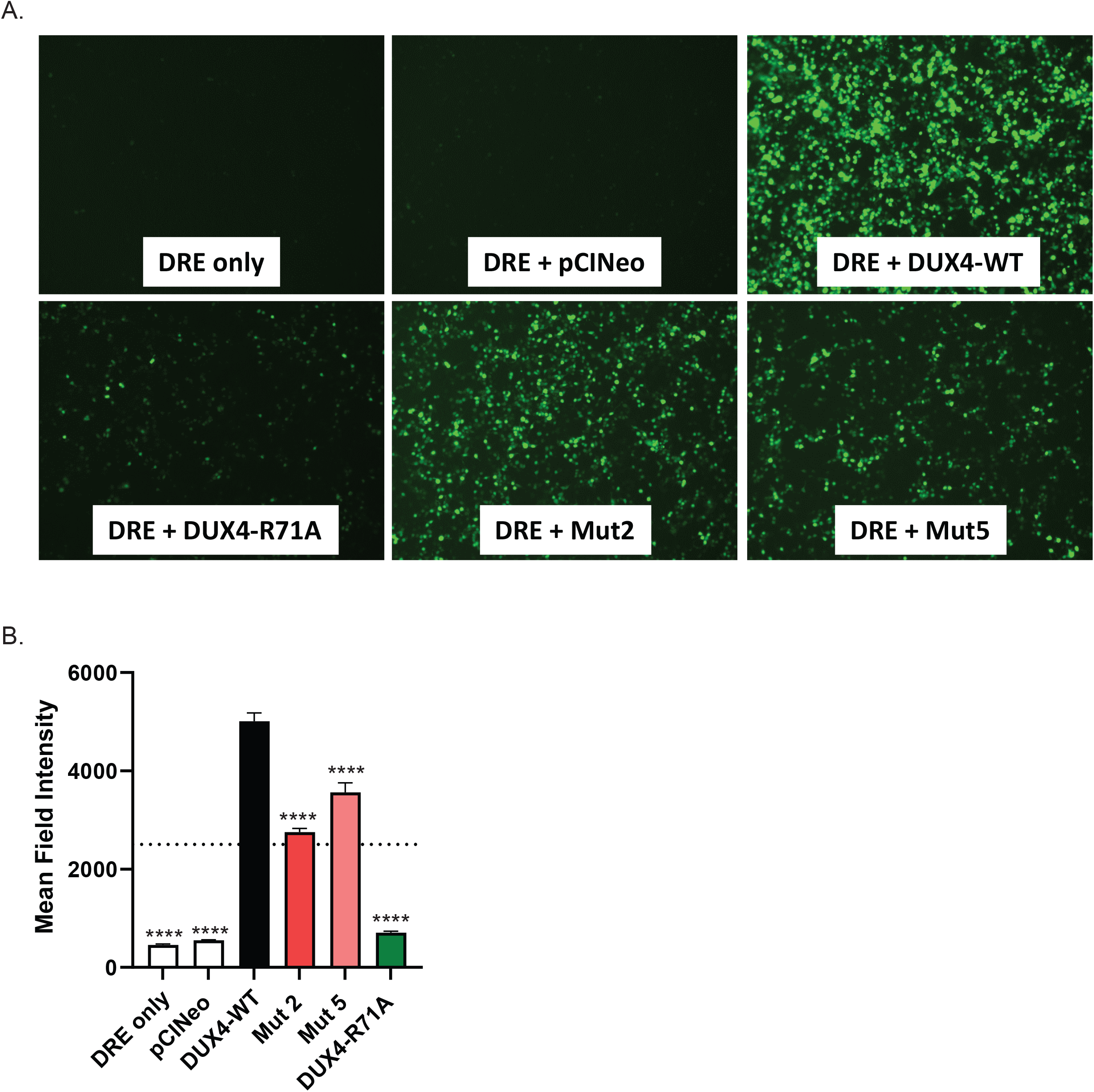
DUX4 PTM mutants have decreased transactivation. HEK293 cells were co-transfected with DUX4-activated fluorescence reporter (DRE) and wild type DUX4, DUX4 R71A or empty vector and GFP expression was visualized 24 hours later. (A) Representative fluorescence photomicrograph of HEK293 cells transfected with the indicated plasmids and DUX4-responsive GFP reporter. (B) Quantification of DUX4-activated fluorescence reporter. N=3 experiments. Asterisk (*) indicates significant differences compared to DUX4-WT. **** P<0.0001. One-way ANOVA test. Dashed line indicates the value which is 50% of DUX4-WT.

**FIGURE 4.**
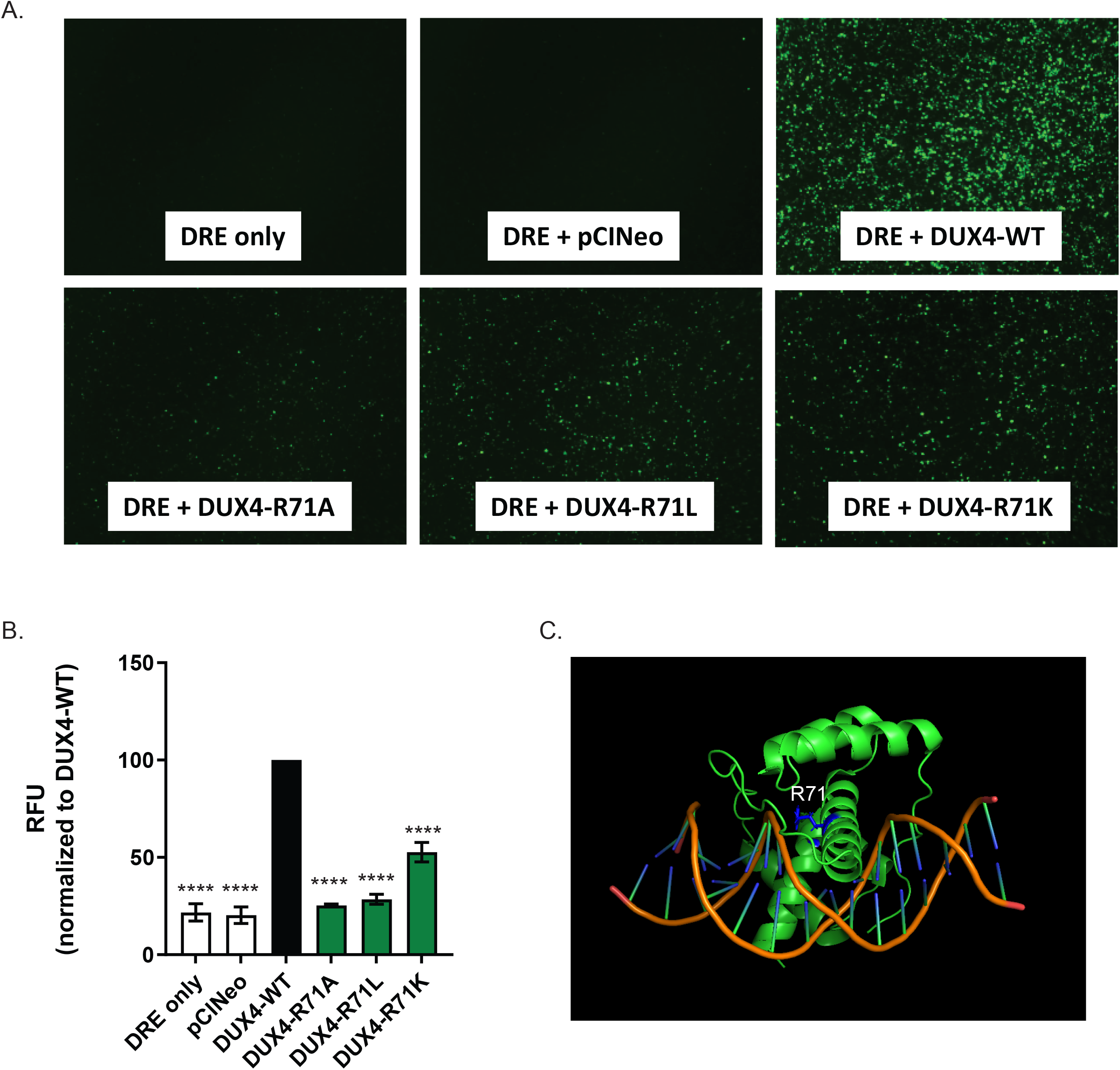
DUX4-R71 mutants have a gradient of decreased transactivation. HEK293 cells were co-transfected with DUX4-activated fluorescence reporter (DRE) and wild type DUX4, DUX4 R71 mutants, or empty vector (pClneo) and GFP expression was visualized 24 hours later. (A) Representative fluorescence photomicrograph of HEK293 cells transfected with the indicated plasmids and DUX4-responsive GFP reporter. (B) Quantification of DUX4-activated fluorescence reporter normalized to DUX4-WT. n=3 independent experiments. (C) The localization of R71 in the crystal structure of DUX4 with DNA. Asterisk (*) indicates significant differences compared to DUX4-WT. **** P<0.0001. One-way ANOVA test. RFU: relative fluorescence units.

To this point in the study, we performed all experiments in human embryonic kidney cells (HEK293). Since FSHD primarily affects muscle, we next sought to confirm the non-toxic effects of DUX4 mutants in human muscle cells. Although we identified 5 non-toxic DUX4 mutants with reduced ability to transactivate a DUX4-responsive reporter construct, 3 contained mutations in the same R71 amino acid, and we therefore used one (R71A) to represent the group in all subsequent experiments. We transfected human myoblasts with plasmids expressing these 3 PTM mutants and measured cell viability using a Trypan Blue assay. While wild-type DUX4 caused significant myoblast cell death, the 3 DUX4 PTM mutants had no deleterious effects on myoblast viability (Fig 4A). Importantly, these PTM mutants were not associated with decreased DUX4 protein expression or altered nuclear localization when expressed in HEK293 cells (Fig 4 B, C).

We next assessed the ability of DUX4-R71A, DUX4-Mutant 2 and DUX4-Mutant 5 to activate 4 DUX4 target genes, *PRAMEF12, ZSCAN4, TRIM43, and LEUTX*. We expressed wild type or DUX4 R71A, DUX4-S29D/S31D/T106D (Mutant 2) and DUX4-T99E/T102E/S104E/T106E (Mutant 5) in HEK293 cells and human myoblasts, and examined DUX4 target gene expression by quantitative RT-PCR. While wild-type DUX4 activated all four biomarkers in HEK293s and myoblasts, cells transfected with the three DUX4 mutants showed significantly decreased *PRAMEF12, ZSCAN4, TRIM43*, and *LEUTX* expression (Fig 5A, B).

**FIGURE 5.**
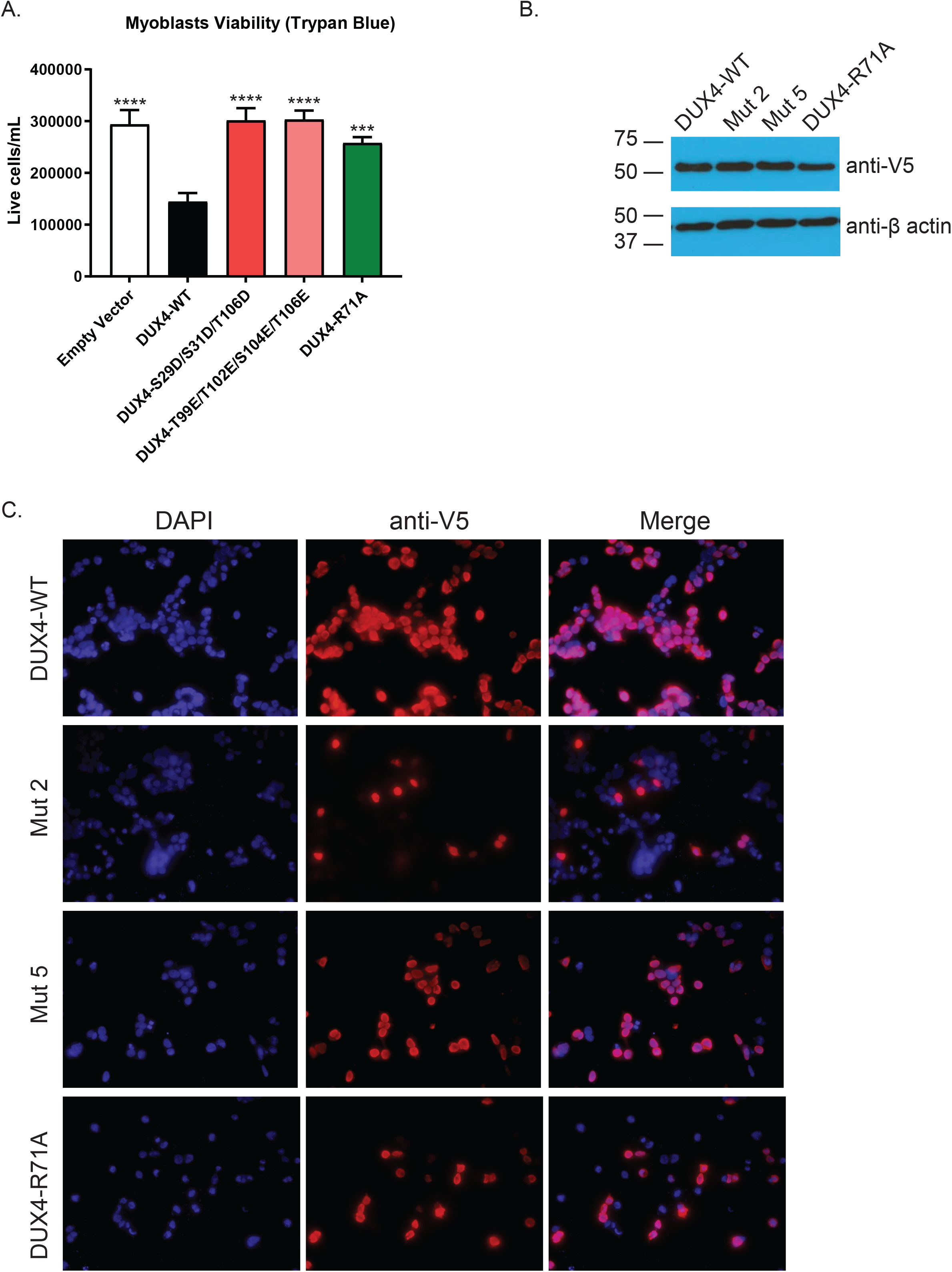
DUX4 PTM mutants protect against cell death in human myoblasts without affecting DUX4 expression or nuclear localization. (A) Human myoblasts were transfected with wild type DUX or the indicated DUX4 PTM mutant and viability was assessed 48 hours later with trypan blue counting. n = 3 independent experiments. Asterisk (*) indicates significant differences compared to DUX4-WT. *** P<0.001, **** P<0.0001. One-way ANOVA test. (B, C) HEK293 cells were transfected with wild type DUX4 or the indicated PTM mutants and expression was assessed by western blotting and immunofluorescence 24hrs later. (B) Western blot demonstrating DUX4 protein in HEK293 cells. Representative data from n=3 independent experiments. (C) DUX4 protein was detected using fluorescent anti-V5 antibody (red) 24 hours post-transfection. Nuclei were DAPI stained (purple). Representative data from n=3 independent experiments.

Our data to this point demonstrated that DUX4 arginine methylation inhibition and enhanced serine/threonine phosphorylation on specific amino acids was associated with decreased apoptosis and DUX4 target gene activation. Thus, because DUX4 function could be affected by arginine methylation and serine/threonine phosphorylation, we next sought to identify kinases and arginine methyltransferases that were capable of modifying the DUX4 protein. First, we conducted a comprehensive screen to identify serine/threonine kinases that could directly phosphorylate DUX4. To do this, we performed an *in vitro* radiometric filter binding assay using purified recombinant DUX4 protein and 245 kinases (Fig 6A, B and Supp Table 2). We identified 92 kinases with significant DUX4 phosphorylation activity (activity ratio value greater than 3) (Supp Table 2). We focused subsequent experiments on a subset of the kinases with a high phosphorylation activity (activity ratio of 9 or greater).

**FIGURE 6.**
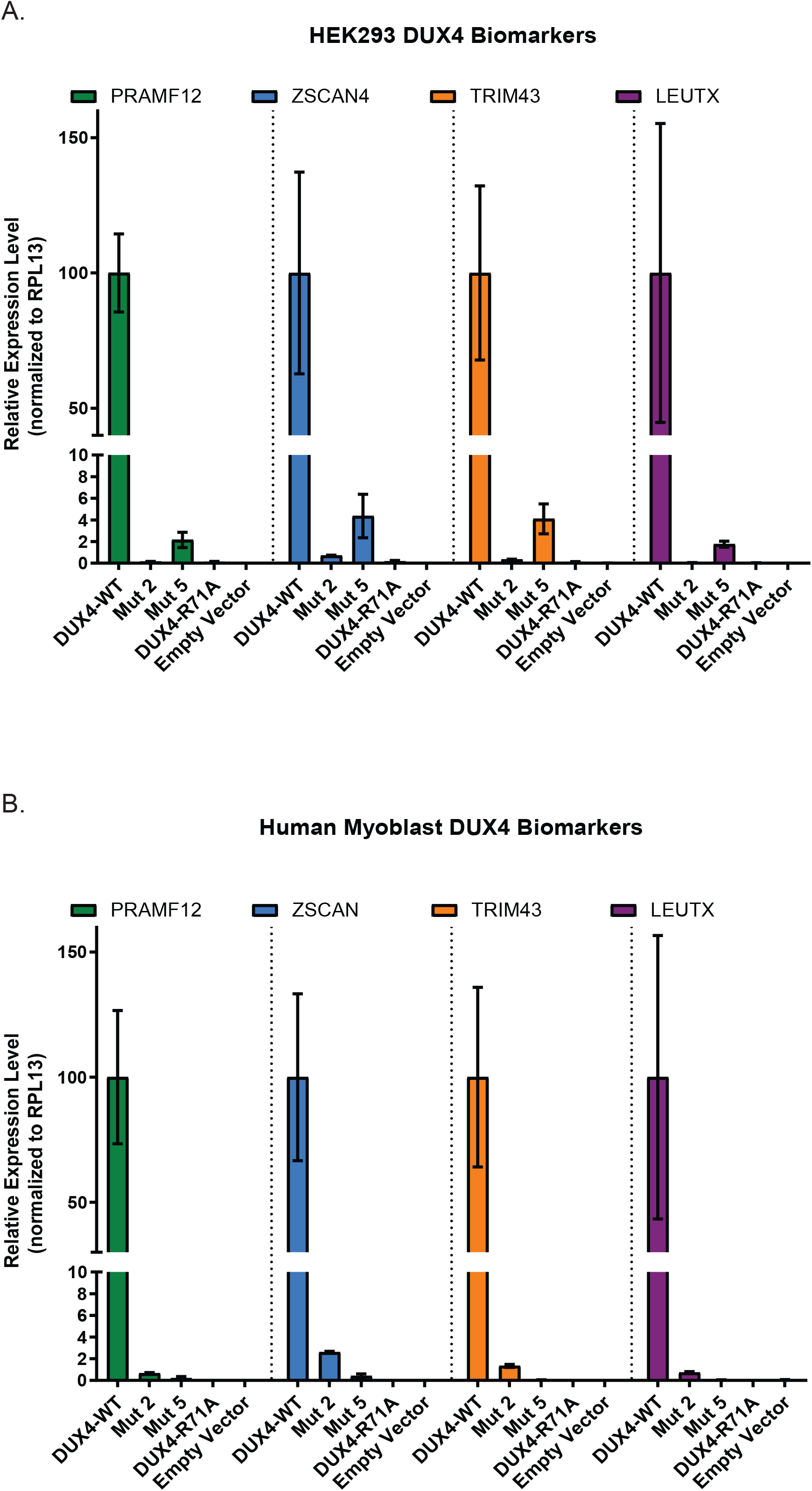
Decreased DUX4 target gene activation in DUX4 PTM mutants. (A) HEK293 cells or (B) myoblasts were transfected with wild type DUX4, empty vector (pClneo), DUX4 R71A, DUX4-Mutant 2 and DUX4-Mutant 5 and quantitative RT-PCR was performed 24 hours later for DUX4 target genes *PRAMF12, ZSCAN4, TRIM43* and *LEUTX*. Gene expression was normalized to the reference gene, RPL13A. n = 3 independent experiments performed in triplicate.

Our finding that DUX4 phosphorylation mimic mutants protect against DUX4-mediated cell death suggested that overexpression of serine/threonine kinases with high DUX4 phosphorylation activity could also rescue the cell death phenotype. We overexpressed DUX4 with a subset of kinases with high activity to DUX4 (Fig 7 C, D) and found the catalytic subunit of protein kinase A, PRKACA, was associated with decreased caspase-3/7 activity (Fig 7E). Another kinase, TBK1, also showed weaker ability to reduce DUX4-associated caspase-3/7 activation *in vitro* (Fig 7D). Importantly, DUX4 and PRKACA form a complex when overexpressed in HEK293 cells (Fig 7F). Taken together, these results demonstrate that DUX4 is a protein kinase A (PKA) substrate and suggested that activation of the PKA pathway could protect against DUX4-mediated toxicity.

**FIGURE 7.**
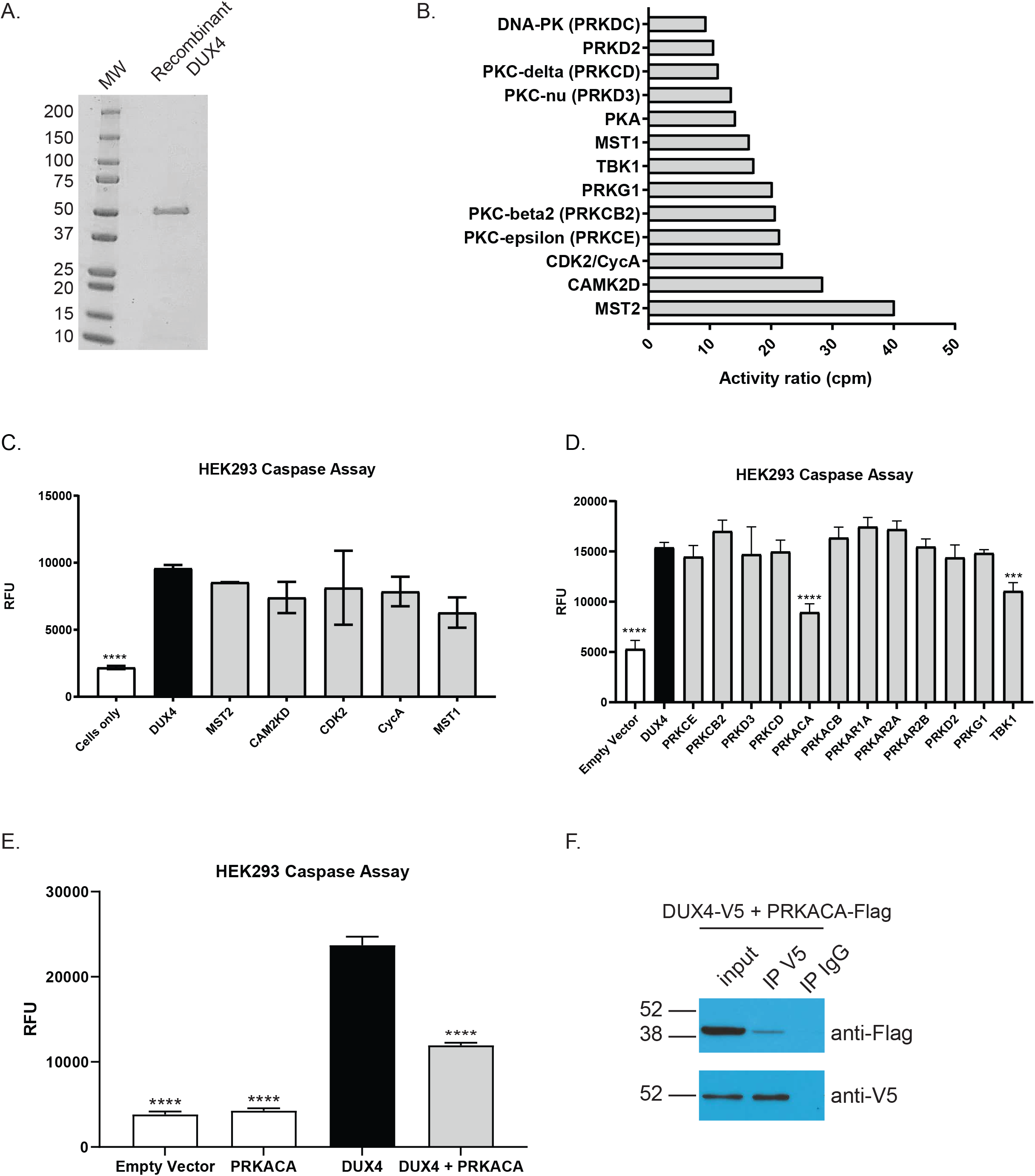
DUX4 is a PKA substrate. A radiometric in vitro kinase assay was performed with (A) purified recombinant DUX4 protein. (B) Activity ratios from a subset of serine/threonine kinases with high activity to DUX4 protein. (C, D) HEK293 cells were transfected with wild type DUX4 and the indicated serine/threonine kinase, and the caspase 3/7 assay was performed 48 hrs later. (E) HEK293 cells were transfected with wild type DUX4 and/or the catalytic subunit of PKA, PRKACA, and the caspase 3/7 assay was performed 48 hours later. Representative data from n=3 independent experiments performed in (C) duplicate and (D, E) triplicate. Asterisk (*) indicates significant differences compared to DUX4-WT. *** P<0.001, **** P<0.0001. Dashed line indicates the value which is 50% of DUX4-WT. One-way ANOVA test. RFU = relative fluorescence units.

To isolate arginine methyltransferases that associate with the DUX4 complex, we used Rapid Immunoprecipitation Mass Spectrometry of Endogenous Proteins (RIME), which combines crosslinking, immunoprecipitation and mass spectrometry to identify transient or distant interactions ^26^. We performed RIME in human myoblasts expressing wild type, V5 epitope-tagged DUX4 and identified protein arginine methyltransferase 1 (PRMT1) as a component of the DUX4 complex (Table 4 and Suppl Table 3). Additionally, we found that DUX4 and PRMT1 interact when overexpressed in HEK293 cells (Fig 8A).

**TABLE 4.** Selected list of DUX4 interacting proteins.

| Protein name | Gene Name | Enzyme Function |
| --- | --- | --- |
| Protein arginine N-methyltransferase 1 | PRMT1 | Methylation |
| rRNA 2'-O-methyltransferase fibrillarin | FBL | Methylation |
| DNA-dependent protein kinase catalytic subunit | PRKDC | Phosphorylation |
| Histone deacetylase 1 | HDAC1 | Acetylation |
| Histone deacetylase 2 | HDAC2 | Acetylation |
| Ubiquitin-like modifier-activating enzyme 1 | UBA1 | Ubiquitination |
| Dolichyl-diphosphooligosaccharide--protein glycosyltransferase subunit 1 | RPN1 | Glycosylation |

Modifying enzymes identified by RIME are listed with enzymatic function.

**FIGURE 8.**
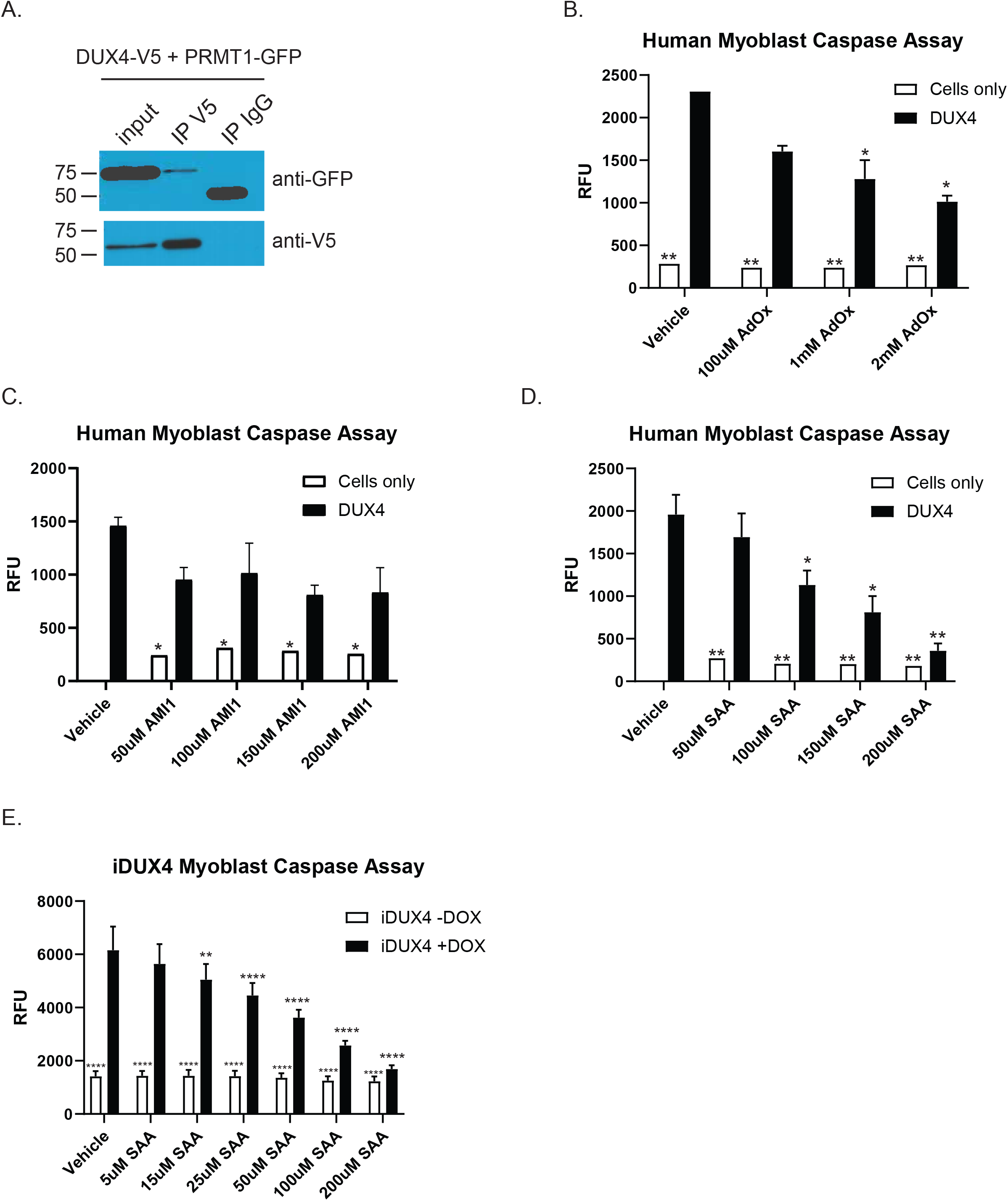
Arginine methylation inhibitors protect against DUX4-mediated cell death in human myoblasts. (A) Co-immunoprecipitation of DUX4 and PRMT1 in HEK293 cells. Representative data from n = 3 independent experiments. Human myoblasts were transfected with DUX4 or no DNA in the presence or absence of increasing concentrations of (B) AdOx, (C) AMI1 or (D) SAA and the caspase 3/7 assay was performed 48 hours later. Representative data from n=3 independent experiments performed in duplicate or singlet. (E) iDUX4 human myoblasts were treated with increasing concentrations of SAA and the caspase 3/7 assay was performed 48 hours later. n = 4 independent experiments performed in triplicate. Asterisk indicates significant differences compared to DUX4-WT. * P<0.05, ** P<0.01, **** P<0.0001. One-way ANOVA test. RFU: relative fluorescence units.

The observation that a DUX4 arginine methylation null mutant protects against cell death and that DUX4 interacts with an arginine methyltransferase raises the possibility that arginine methylation inhibitors could be protective in FSHD disease models. To investigate this, we tested three compounds with activity against PRMTs, the global methylation inhibitor adenosine dialdehyde (AdOx) ^27^, type 1 PRMT inhibitor arginine methyltransferase inhibitor 1 (AMI1) ^28^ and PRMT1 inhibitor salvianolic acid A (SA-A) ^29^. We performed caspase-3/7 assay in human myoblasts expressing DUX4 in the presence or absence of AdOx, AMI1, or SA-A. We found that SA-A and AdOx led to significantly decreased caspase cleavage in a dose-responsive manner (Figure 8B,D), while AMI1 did not (Fig 8C). SA-A was also protective in the inducible DUX4 myoblast cell line (iDUX4) ^25^ (Fig 8E). We conclude that targeting arginine methylation protects against DUX4-mediated cell death in human myoblasts.

## DISCUSSION

This is the first study to identify methylation and phosphorylation as critical regulators of DUX4-mediated toxicity. We characterized arginine methylation null mutants and two phosphorylation mimic mutants that prevented DUX4-associated apoptosis and decreased DUX4 target gene activation. DUX4 forms a complex with arginine methyltransferase PRMT1 and serine/threonine kinase PKA. Importantly, arginine methylation inhibitors protect against DUX4-mediated toxicity in human myoblasts. Taken together, these results suggest that DUX4 PTMs may be druggable targets for FSHD therapy.

Arginine methylation plays an important role in the regulation of transcription factors by affecting DNA binding and target gene activation ^30^. Consistent with this, we found that DUX4 R71 is critical for DUX4 transactivation activity. We demonstrate that the arginine methyltransferase PRMT1 is a component of the DUX4 complex. PRMT1 is the predominant type I arginine methyltransferase accounting for 85% of arginine methyltransferase activity in mammalian cells ^31,32^. In skeletal muscle, PRMT1 plays a central role in differentiation and regeneration ^33,34^. In particular, PRMT1 methylation of the transcription factor MyoD affects DNA binding and target gene activation in C2C12 cells^50^ and PRMT1 inhibition impacts the recruitment of the transcription factor Myc to specific target promoters ^35^. Further studies are required to determine whether PRMT1 is directly responsible for recruiting DUX4 to target gene promoters and the molecular consequences of the DUX4-PRMT1 interaction.

Our data demonstrate that serine/threonine phosphorylation of DUX4 and overexpression of DUX4 with the catalytic subunit of serine/threonine kinase PKA protect against DUX4-mediated toxicity. These results reveal that serine/threonine phosphorylation of the DUX4 protein is protective. The beneficial effects of DUX4 phosphorylation could result from diminished activation of target genes or by affecting protein recruitment to the DUX4 transcriptional complex resulting in impaired transactivation. There is evidence for such mechanisms in other systems. For example, in the silkworm, PKA phosphorylation of the transcription factor BR-C impairs transcriptional activity^53^ while a phosphorylation mimic mutant of EZH2, a PKA substrate, has enhanced interactions with STAT3, resulting in decreased oncogenic activity ^36^,^37^. Interestingly, activation of the PKA pathway represses *DUX4* mRNA expression ^38^. These results suggest a dual mechanism by which the PKA pathway negatively regulates DUX4 by repressing DUX4 expression at the mRNA level and abrogating DUX4-mediated toxicity at the protein level.

Enhancing DUX4 serine/threonine phosphorylation by activating the PKA pathway or inhibiting serine/threonine phosphatases could serve as novel therapeutic approaches for FSHD. In a prior study, PKA pathway activation with beta-adrenergic receptor agonists, cAMP analogs and a constitutively active PKA mutant led to decreased *DUX4* mRNA expression and decreased DUX4 target gene activation in FSHD patient myotubes ^38^. It remains to be determined how PKA pathway agonists affect the DUX4 protein. Serine/threonine phosphatases PP2A and calcineurin antagonize PKA ^39,40^. Our work suggests that inhibitors of PP2A and calcineurin could counteract the dephosphorylation of DUX4, resulting in protection from DUX4-mediated cell death. Calcineurin inhibitors Cyclosporine A and tacrolimus are FDA approved drugs for immunosuppression ^41^. Treating FSHD with immunosuppressants such as prednisone has not been shown to be efficacious ^42^. Consistent with this, a recent case report of a FSHD patient treated with a 12-week course of tacrolimus and prednisone showed disease progression on this regimen ^43^. Alternatively, LB-100 is a PP2A inhibitor with anti-tumor activity which is safe in adult patients with progressive solid tumors ^44^. LB-100 is currently being investigated in clinical trials for cancer therapy (NCT03027388, NCT04560972). Future studies are necessary to determine whether serine/threonine phosphatase inhibitors such as LB-100 maintain the protective phosphorylation status of the DUX4 protein and improve DUX4-mediated toxicity.

While PRMT inhibitors have been studied extensively in cancer with several compounds advancing to clinical trials ^22^, this is the first report to demonstrate a therapeutic role for PRMT inhibitors in muscle disease. We have identified a promising compound, SA-A, which rescues DUX4-mediated cell death in human myoblasts. SA-A is derived from the root of *Salvia miltiorrhiza* which has been used in traditional medicine in Asian countries for a wide range of ailments including heart disease, dementia and cancer ^45^. Oral administration of SA-A is safe in rodents ^46,47^ and there is a planned clinical trial for SA-A in patients with arterial ischemic stroke (NCT04931628). SA-A inhibits PRMT1 activity *in vitro* ^29^, however it also protects against oxidative stress ^47,48^. DUX4 expression in proliferating myoblasts is associated with increased susceptibility to oxidative stress and suppression of the glutathione redox pathway ^8,49^. Therefore, SA-A could exert its protective effects in FSHD disease models by inhibiting PRMT1 and mitigating oxidative stress.

In conclusion, in this first study to explore DUX4 post-translational modification, we identified and functionally characterized several modified amino acids in the DUX4 protein. We found several DUX4 residues that are essential for DUX4 target gene activation and DUX4-mediated cell death. In addition, we identified potential modifiers of the DUX4 transcriptional complex including PRMT1 and PKA, which may serve as novel therapeutic targets for FSHD. This work lays the foundation for future mechanistic and translational studies to explore the therapeutic potential of targeting DUX4 PTMs in FSHD disease models *in vitro* and *in vivo*.

## TABLES

**Supplemental Table 1.**
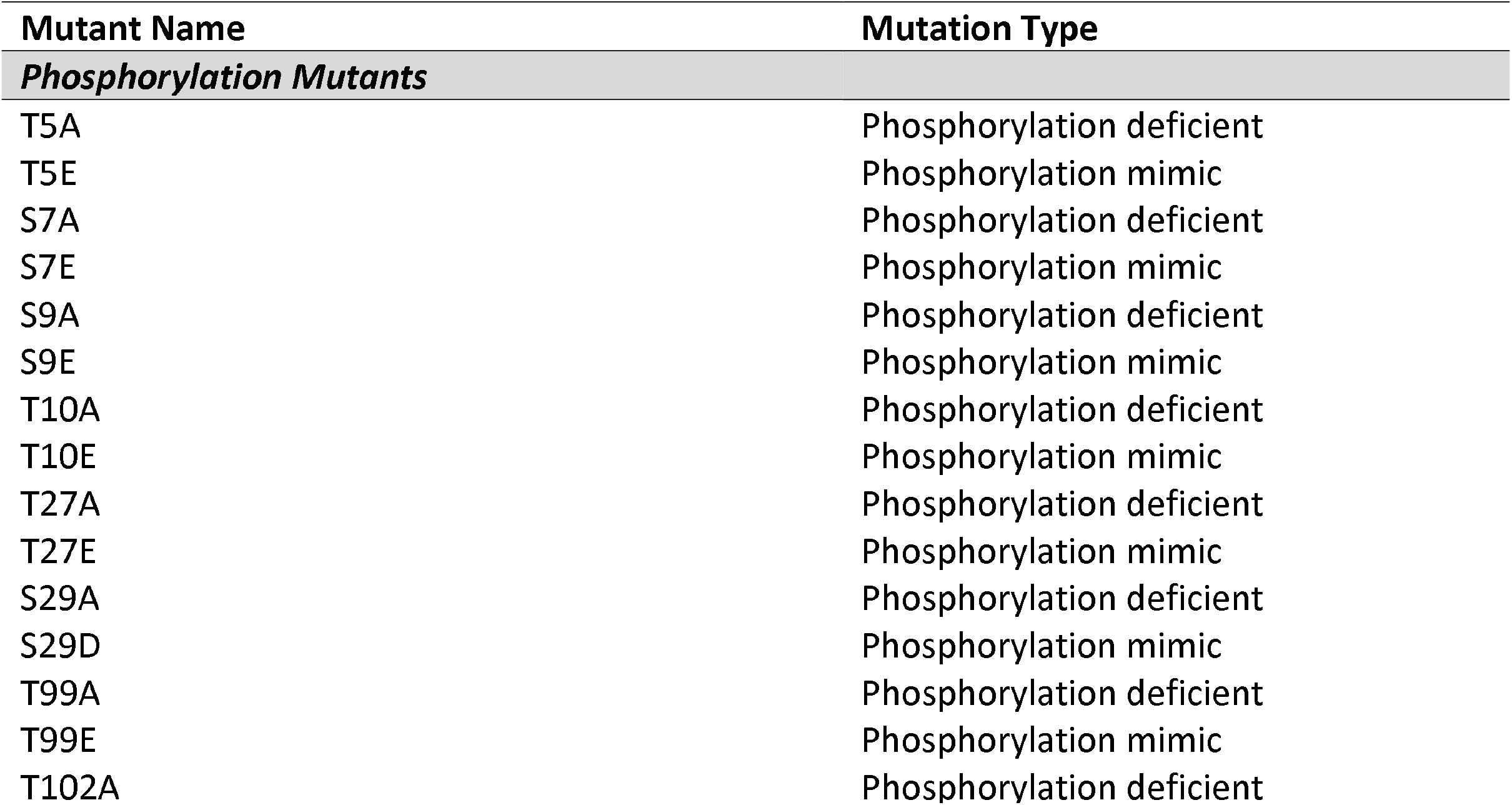
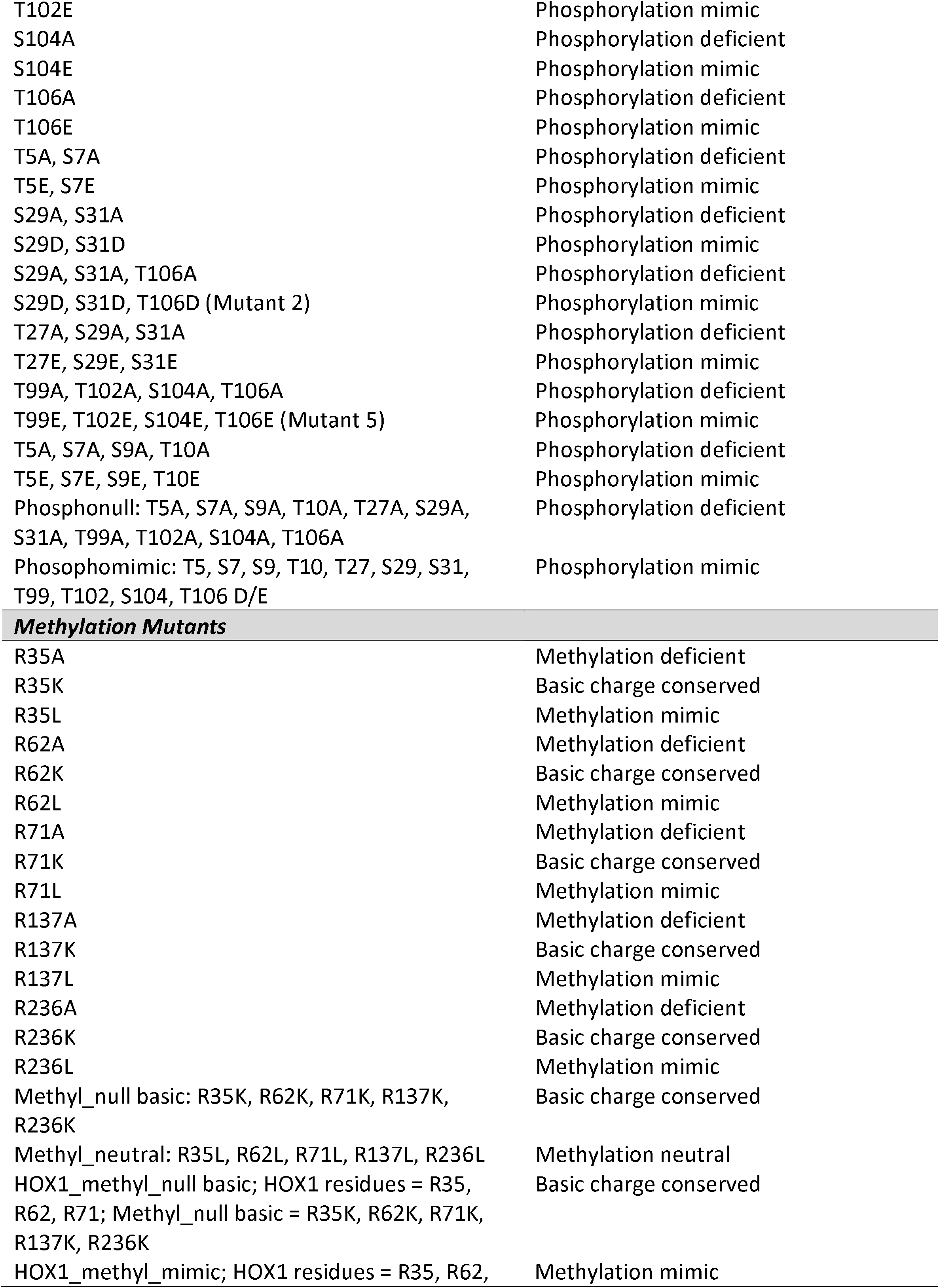
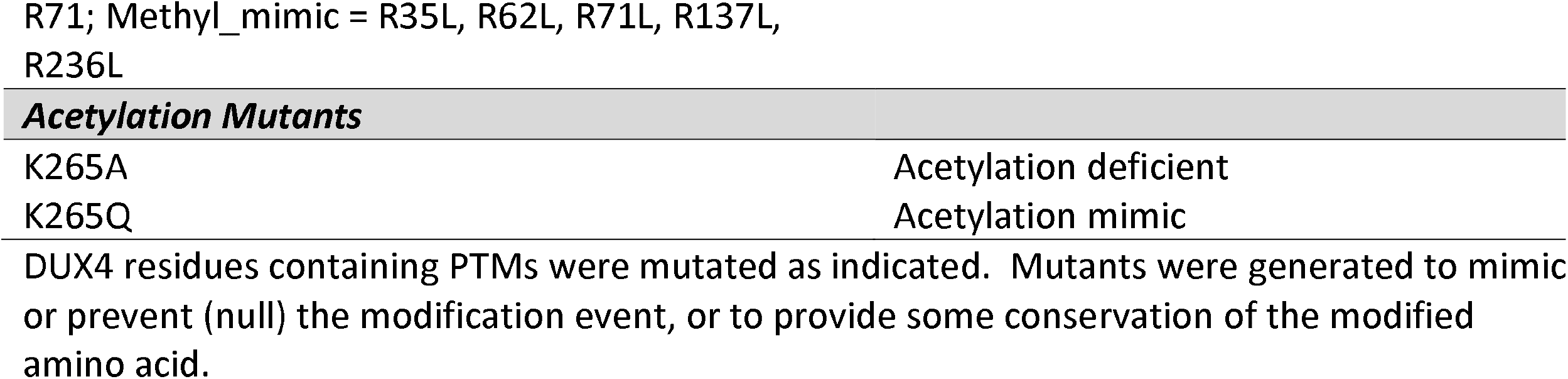
DUX4 Mutagenesis Strategy.

**Supplemental Table 2. Phosphorylation profile of DUX4 with serine/threonine kinases**.A radiometric protein kinase filter-binding assay was used for measuring the kinase activity of 245 serine/threonine kinases. The activity value (raw counts of the kinase assay as measured in the filter plate assay), the normalized kinase autophosphorylation value, the median of three background values of the sample protein and the corrected activity value (raw value minus sample protein background) are reported in the table. The activity ratio value for each kinase describes the ratio between the activity of the particular kinase with the DUX4 protein and without the DUX4 protein. A ratio value of >3 may be considered as significant.

**Supplemental Table 3. Proteins associated with the DUX4 complex in human myoblasts using the RIME assay**.RIME (Rapid Immunoprecipitation Mass Spectrometry of Endogenous Proteins) was carried out using an antibody against V5 tag and 100ug of chromatin from DUX4.V5 transfected human myoblasts to identify proteins that interact with DUX4 using mass spectrometry. The file includes a summary of the proteins enriched in the RIME analysis. Two independent experiments were performed (R1 and R2). The enriched protein list contains uniquely identified proteins for all samples after removing proteins present in the IgG negative control. Three lists were generated – two corresponding to the proteins identified uniquely in one of the two replicates, and one corresponding to proteins identified in both replicates. The file also includes a list of total spectrum counts for all proteins and peptides identified in all samples.

## Acknowledgements

This study was funded by grants from The Chris Carrino Foundation (S.Q.H. and J.O.E); Friends of FSHD Research (J.O.E., S.C. and N.Y.S.); The FSHD Society (J.O.E., A.R., and N.Y.S.); National Institute of Arthritis and Musculoskeletal and Skin Diseases Center of Research Translation in Muscular Dystrophy Therapeutic Development (1P50AR070604-01, S.Q.H.); The National Institute of Neurological Disorders and Stroke Translational R21/R23 (R21NS101166, S.Q.H.); The National Institute of Arthritis and Musculoskeletal and Skin Diseases (1RO1AR062123-05, S.Q.H.); The University of Massachusetts Medical School Wellstone Center for FSHD (5U54HD060848-10, S.Q.H.); National Institutes of Health (S10 OD018056, M.A.F.); American Academy of Neurology, Muscle Study Group and American Brain Foundation Clinical Research Training Scholarship in Neuromuscular Disease (R.N.K.)

We thank Dr. Mark Bedford, MD Anderson Cancer Center, and Dr. Jocelyn Cote, University of Ottawa, for generously providing the GFP-PRMT1 constructs.

## Author Contributions

R.N.K, J.O.E., M.A.F., and S.Q.H. were responsible for the conception and design of the study. R.N.K, J.O.E., L.W., S.C., A.R., N.Y.S., M.E.H., L.Z., and O.E.B. were responsible for acquisition and analysis of data. R.N.K., J.O.E., M.A.F., and S.Q.H. were responsible for drafting the text and preparing the figures.

